# WOX9 functions antagonistic to STF and LAM1 to regulate leaf blade expansion in *Medicago truncatula* and *Nicotiana sylvestris*

**DOI:** 10.1101/2020.03.23.003715

**Authors:** Tezera W. Wolabu, Hui Wang, Dimiru Tadesse, Fei Zhang, Marjan Behzadirad, Varvara E. Tvorogova, Haggagi Abdelmageed, Ye Liu, Naichong Chen, Jianghua Chen, Randy D. Allen, Million Tadege

## Abstract

Plant specific WOX family transcription factors are known to regulate embryogenesis, meristem maintenance and lateral organ development. Modern clade WOX genes function through a transcriptional repression mechanism, and the intermediate clade transcriptional activator WOX9 functions with the repressor WOX genes in embryogenesis and meristems maintenance, but the mechanism of this interaction is unclear. *WOX1* homologues *STF* and *LAM1* are required for leaf blade outgrowth in *Medicago truncatula* and *Nicotiana Sylvestris*, respectively. Here we show that *WOX9* negatively regulates leaf blade outgrowth and functions antagonistically to *STF* and *LAM1*. While *NsWOX9* ectopic expression enhances the *lam1* mutant phenotype, and antisense expression partially rescues the *lam1* mutant, both overexpression of *NsWOX9* and knockout by CRISPR/Cas9 genome editing in *N. sylvestris* resulted in a range of severe leaf blade distortions, indicating that controlled negative regulation by NsWOX9 is required for proper blade development. Our results indicate that direct repression of WOX9 transcriptional activation activity by the transcriptional repressor STF/LAM1 is required for correct blade architecture and patterning in *M. truncatula* and *N. sylvestris*. These findings suggest that a balance between transcriptional activation and repression mechanisms by direct interaction of activator and repressor WOX genes may be required for cell proliferation and differentiation homeostasis, and could be an evolutionarily conserved mechanism for the development of complex and diverse morphology in higher plants.

**One sentence summary:** WOX9 negatively regulates blade outgrowth antagonizing STF function but directly repressed by STF indicating WOX-mediated homeostasis in cell proliferation and differentiation during leaf morphogenesis.

## Introduction

WUSCHEL-related homeobox (WOX) factors are plant-specific transcriptional regulator proteins that contain a DNA binding homeodomain similar to WUSCHEL (WUS), the founding member of the family from Arabidopsis. Several elegant studies demonstrated that the WOX family is involved in the regulation of a wide range of key developmental programs ranging from the modulation of zygotic development and embryogenesis by WOX2, WOX8, and WOX9 (Haecker et al., 2004; Breuninger et al., 2008; Ueda et al., 2011) to maintenance of shoot and root apical meristems orchestrated by WUS and WOX5, respectively (Mayer et al., 1998; Sarkar, 2007), along with several other developmental pathways (Matsumoto and Okada, 2001; Park et al., 2005; Deyhle et al., 2007; Shimizu et al., 2009; Vandenbussche et al., 2009; Hirakawa et al., 2010; Ji et al., 2010; Tadege et al., 2011b; Nakata et al., 2012).

*WUS* and its homologues in other species, including *TERMINATOR* (*TER*) in petunia, *ROSULATA (ROA)* in Antirrhinum, and *HEADLESS* (*HDL*) in Medicago, are required for shoot apical meristem (SAM) maintenance (Laux et al., 1996; Mayer et al., 1998; Stuurman et al., 2002; Kieffer et al., 2006; Meng et al., 2019; Wang H, 2019). Loss of *WUS* function in the *wus-1* Arabidopsis mutant results in premature termination and arrest of the SAM and floral meristem, but the SAM re-establishes itself to resume growth while the process repeats itself, leading to altered plant morphology (Laux et al., 1996; Mayer et al., 1998). In the *hdl* mutant of *M. truncatula*, termination of SAM and axillary meristems is permanent. Since the SAM fails to re-establish itself, these plants only make leaves throughout development (Tadege et al., 2015; Meng et al., 2019; Wang H, 2019). The *hdl* mutant also shows altered leaf shape (Meng et al., 2019; Wang et al., 2019), which was not detected in the *wus* mutant. However, the *wus wox1 prs* triple mutant showed a stronger leaf phenotype than the *wox1 prs* double mutant (Zhang F, 2015), suggesting that WUS may also have a redundant function in leaf development. Although *WUS* transcript is specifically expressed in the organizing center (OC) of the SAM (Mayer et al., 1998), it is likely that a non-cell autonomous signal from WUS may contribute to blade outgrowth, since the WUS protein itself is shown to move from the OC to the stem cell region (Yadav et al., 2011; Daum et al., 2014).

An intimate connection exists between WUS and the phytohormone cytokinin, and this appears to be true with some other WOX orthologs (Tadege and Mysore, 2011; Tadege, 2016; Wang, 2017). While WUS activity is modulated by cytokinin in the SAM and axillary meristem (Wang et al., 2017; Snipes et al., 2018), WUS promotes cytokinin activity in the shoot stem cell niche by repressing type-A *ARABIDOPSIS RESPONSE REGULATOR (ARR)* genes *ARR5, ARR6, ARR7*, and *ARR15* (Leibfried et al., 2005) to activate cell proliferation, and WUS physically interacts with the transcriptional co-repressor TOPLESS (TPL) to repress target genes that promote cell differentiation (Kieffer et al., 2006; Causier et al., 2012; Yadav et al., 2013). Genes that encode polarity factors which impart adaxial or abaxial identity and maintenance of a differentiated state to the leaf blade tissues are among the targets directly repressed by WUS (Yadav et al., 2013). This uncovers a mechanism by which WUS maintains undifferentiated stem cells in the SAM, in addition to the well-established CLE peptide signaling (Schoof et al., 2000; Somssich et al., 2016; Hu C, 2018). Although WUS is reported to be a bifunctional transcription factor exhibiting both transcriptional repression and activation activities (Ikeda et al., 2009), its SAM maintenance activity and interaction with cytokinin are shown to be linked to its WUS box (Ikeda et al., 2009; Dolzblasz et al., 2016; Snipes et al., 2018), suggesting that WUS primarily functions as a transcriptional repressor. Interestingly, *WUS* is reported to be activated by WOX9/STIP, which is also required for shoot meristem maintenance (Wu et al., 2005) and embryo development (Wu et al., 2007; Ueda et al., 2011). However, WOX9 is reported to be a strong transcriptional activator (Lin et al., 2013), and it is unclear whether WUS and WOX9 employ the same mechanism in shoot meristem maintenance.

The WOX9 gain-of-function *sitp-D* mutant displays wavy leaf margins indicating problems with cell division in leaf primordium (Wu et al., 2005). But, the *stip* loss-of-function mutant is arrested at the seedling stage (Wu et al., 2005), and it is unclear if WOX9/STIP plays a specific role in leaf blade development. However, WOX function in leaf development is not uncommon in Arabidopsis and several other species. WOX1 and PRS/WOX3 in Arabidopsis (Vandenbussche et al., 2009; Nakata et al., 2012) and their homologues in maize, rice, petunia, *Medicago* and woodland tobacco regulate leaf blade development (Nardmann et al., 2004; Vandenbussche et al., 2009; Tadege et al., 2011b; Zhuang et al., 2012Cho et al., 2013; Ishiwata et al., 2013). Unlike leaf polarity factors that are adaxial or abaxial-specific (Waites et al., 1998; Sawa, 1999; Siegfried, 1999; Kerstetter et al., 2001; McConnell et al., 2001; Iwakawa et al., 2002), the WOX genes *STF*, *WOX1* and *PRS*, are expressed in the middle at the adaxial-abaxial juxtaposition to control medial-lateral outgrowth of the leaf blade (Tadege et al., 2011a,b; Nakata and Okada, 2012; Nakata et al., 2012), suggesting a novel mechanism for blade expansion.

*M. truncatula* STF or *N. sylvestris* LAM1 is a transcriptional repressor (Lin, 2013; Lin et al., 2013) and its repression activity is conferred by its WUS box and STF box motifs (Zhang et al., 2014; Zhang and Tadege, 2015). The DNA binding STF homeodomain (HD) and the repression motifs (WUS box and STF box) are critically required for blade outgrowth function (Lin et al., 2013; Zhang et al., 2014; Zhang et al., 2019). Interestingly, all of the WUS clade Arabidopsis WOX transcription factors (WUS and WOX1-WOX7), which have transcriptional repression activity, can substitute for LAM1 function (Lin et al., 2013), suggesting that modern/WUS clade WOX members have a conserved transcriptional repression mechanism in meristem maintenance and lateral organ development, with specificity conferred by cis elements that drive specific expression patterns.

Intermediate clade WOX members have intact HD but lack repression domains. Here we show that homologues of the intermediate clade transcriptional activator WOX9, namely, MtWOX9-1, MtWOX9-2, and NsWOX9, negatively regulate leaf blade outgrowth in *M. truncatula* and *N. sylvestris*. These factors function antagonistically to STF or LAM1, and exacerbate the *lam1* phenotype. Suppression of *NsWOX9* transcript levels by antisense technology partially rescues the *lam1* mutant leaf blade while the introduction of knockout mutations in the native *NsWOX9* gene by multiplex gRNA genome editing severely affected leaf blade symmetry and expansion. Our results suggest that direct and antagonistic interactions between transcriptional repressor and activator *WOX* genes may be important to balance cell proliferation with differentiation in acquiring complex morphology in higher plants.

## Results

### Ectopic expression of *WOX9* enhances *stf* and *lam1* mutant phenotypes

We have previously shown that, while the WUS clade repressor *WOX* genes of Arabidopsis including *WUS* and *WOX1-WOX7* complement the *lam1* mutant phenotype when driven by the *STF* promoter, the intermediate clade *WOX9* expression exacerbates the *lam1* mutant phenotype (Lin et al., 2013). Therefore, we decided to investigate whether the unique activity of *WOX9* is conserved in *Medicago truncatula* and *Nicotiana sylvestris* and determine its biological significance in leaf development. We isolated orthologous coding sequences for *AtWOX9* from *M. truncatula (MtWOX9)* and *N. sylvestris (NsWOX9)* and created expression constructs that placed these sequences under control of the *STF* promoter to ectopically express these genes in the *stf* and *lam1* mutant plants. *stf* and *lam1* are severe leaf blade mutants in *M. truncatula* and *N. sylvestris*, respectively, caused by mutation of the *WOX1* orthologs *STF* and *LAM1* (Figures 1B and 1E). The *M. truncatula* genome contains two *WOX9*-like sequences here designated as *MtWOX9-1* and *MtWOX9-2* (Figure S1 and S2). We introduced *MtWOX9-1* driven by the *STF* promoter (*STF::MtWOX9-1*) first into the *stf M. truncatula* plants and eight independent transgenic lines were generated. Expression of *MtWOX9-1* in the *stf* mutant background was confirmed by RT-PCR assays. All of these transgenic lines displayed strongly enhanced mutant phenotype with much narrower leaves and thinner stems compared to the *stf* mutant (Figures 1A to 1C). In addition, leaves and stems were significantly shorter in length, leading to a dwarf phenotype that was not characteristic of the *stf* mutant phenotype (Figures 1B and 1C). Similarly, introduction of this construct into the *lam1* mutant background severely affected both leaf length and width, exacerbating the *lam1* mutant phenotype (Figures 1D to 1F). These results indicate that both the *stf* and *lam1* mutants respond similarly to the activity of *MtWOX9-1*, consistent with the effect of *AtWOX9* ectopic expression in the *lam1* mutant (Lin et al., 2013). We also transformed *35S::MtWOX9-1, 35S::MtWOX9-2* and *35S::NsWOX9* into the *lam1* mutant and obtained severely enhanced mutant phenotypes similar to the *STF::MtWOX9-1* expressing *lam1* lines (Figure 1G to 1I), indicating that these three genes have similar effects on leaf blade outgrowth. In most cases, these transgenic leaves displayed approximately five-fold reduction in leaf length, but this effect appeared to be dependent on the level of *WOX9* transgene expression since plants with high level of transgene expression showed more severe phenotypes compared to *lam1* plants with low level of transgene expression (Supplemental Figure 3).

**Figure. 1.**
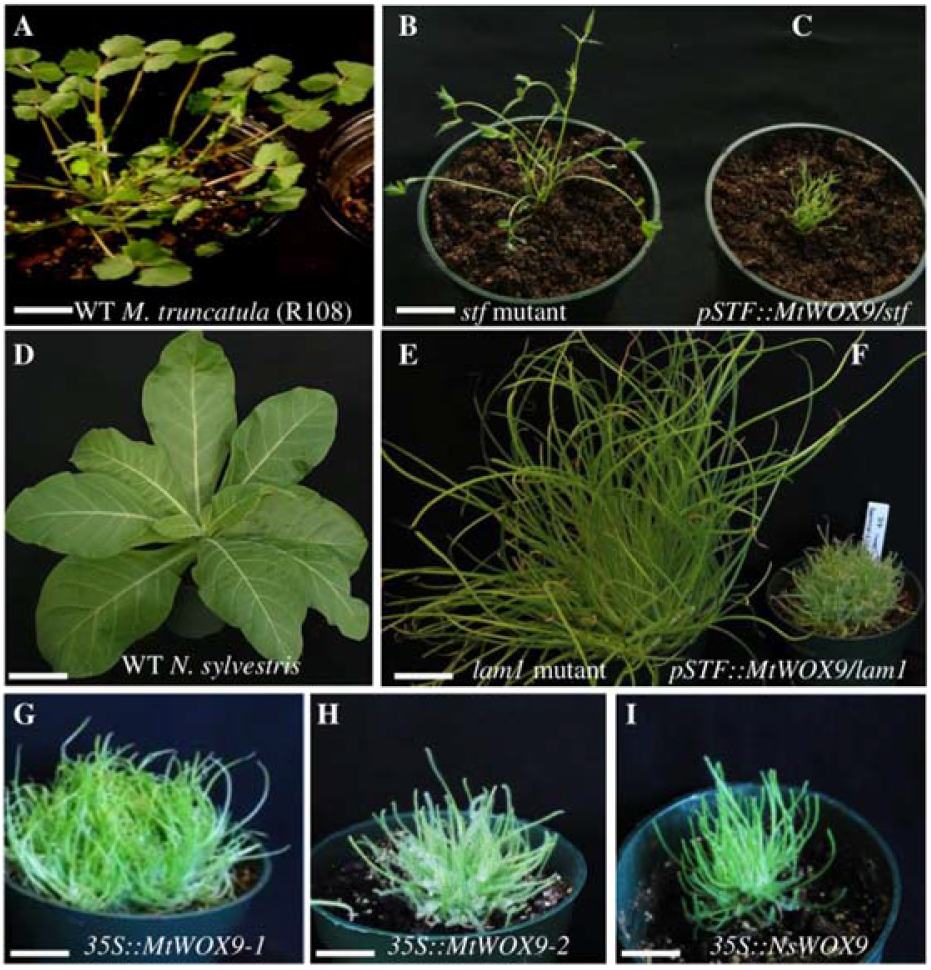
Ectopic expression of *WOX9* enhances *stf* and *lam1* mutant phenotypes. **(A)** Untransformed *M. truncatula* wild type (WT) (R108) plant. **(B)** Phenotype of untransformed *stf* mutant. **(C)** *stf* mutant transformed with *STF::MtWOX9-1*. **(D)** Untransformed *N. sylvestris* WT plant. **(E)** Phenotype of untransformed *lam1* mutant. **(F)** *lam1* mutant transformed with *STF::MtWOX9-1* **(G)** *lam1* mutant transformed with *35S::MtWOX9-1* **(H)** *lam1* mutant transformed with *35S::MtWOX9-2*. **(I)** *lam1* mutant transformed with *35S::NsWOX9*. Plants were 10-weeks (E and F) or 5-weeks old (all the rest). Scale bars: 10 cm.

The enhancement of *stf* and *lam1* mutant phenotypes associated with WOX9 expression suggested to us that *WOX9* acts in opposition to *STF/LAM1* in leaf blade development. Therefore, we introduced an *NsWOX9-antisense* construct into the *lam1* plants to see if reduced levels of *NsWOX9* transcripts could alleviate the mutant phenotype. Indeed, expression of an *NsWOX9-antisense* construct had the opposite effect of *NsWOX9* overexpression, partially rescuing the *lam1* mutant leaf phenotype (Figure 2). However, this partial complementation was limited, and the antisense plants still appeared bushy and failed to make stems. Nonetheless, unlike the untransformed *lam1* mutant control, the *NsWOX9-antisense* leaves showed distinct petioles and blades especially at early stages of development (Figures 2A and 2 B), and the blades were variously branched and curled resulting in unusual leaf structure at maturity (Figures 2C to 2E). These leaf phenotypes suggest that blade outgrowth initiation has significantly progressed in the *NsWOX9-antisense* plants but perhaps aborted before completion.

**Figure. 2.**
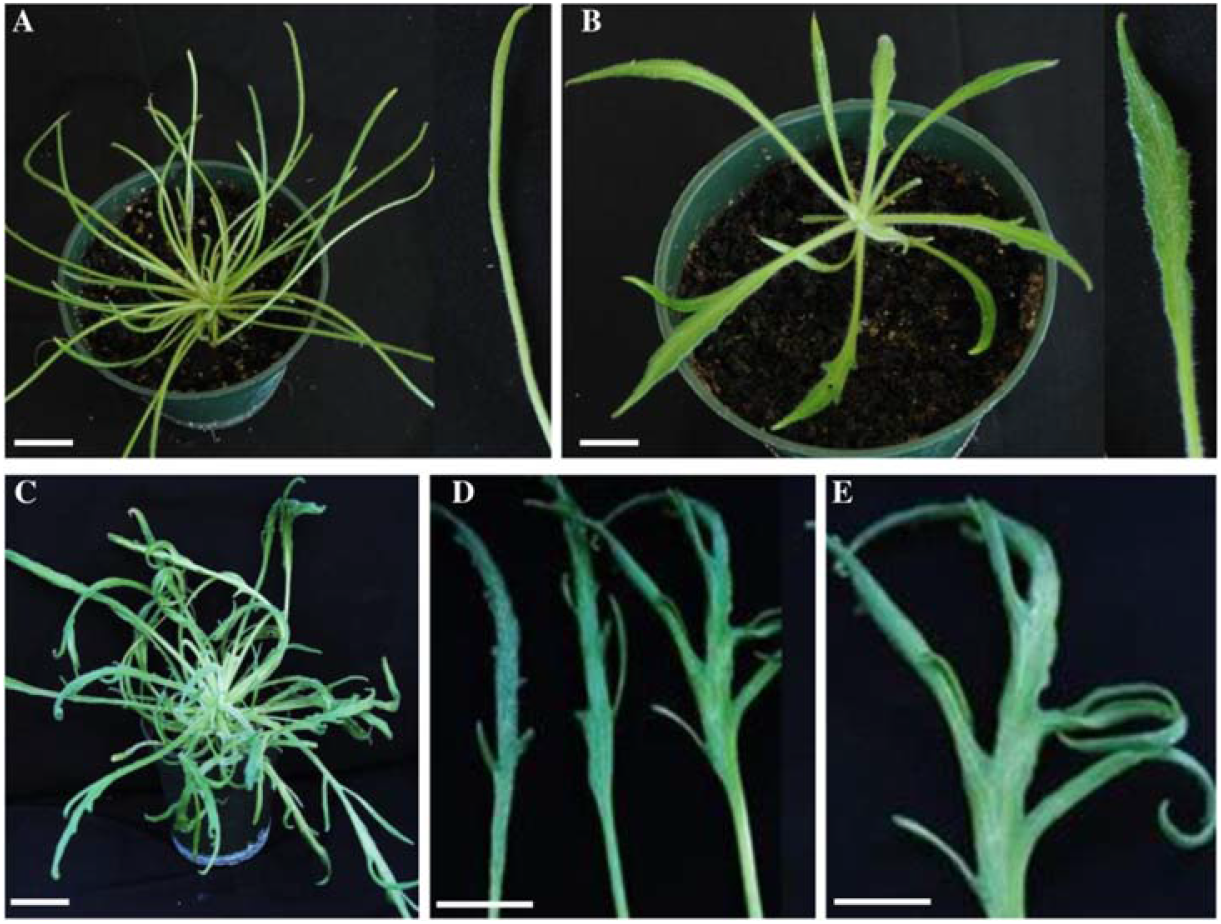
*NsWOX9-antisense* partially rescued *lam1* mutant phenotype. **(A)** Phenotype of *lam1* mutant transformed with *35S::GUS* as control at three weeks of age. Inset on the right is detached leaf close up. **(B)** Partially complemented *lam1* phenotype transformed with *35S::NsWOX9-antisense* construct at three weeks, the inset is close up of a partially complemented leaf blade. Inset on the right is detached leaf close up. **(C)** Partially complemented *lam1* phenotype transformed with *35S::NsWOX9-antisense* construct at seven weeks. **(D)** Representative individual leaves from *35S::NsWOX9-antisense/lam1* plants. **(E)** A magnified view of a leaf in (D). Note the branching and curling of leaves especially in older *35S::NsWOX9-antisense/lam1* plants. Scale bars: A-D, 5 cm, E, 1.5 cm.

To more completely evaluate the effects altered WOX9 expression on the *lam1* mutant phenotype, we carried out structural examination of leaf tissues of WT, *lam1* mutant, *WOX9* overexpressing *lam1* and *WOX9* antisense *lam1* plants (Figures 3A to 3D).

**Figure. 3.**
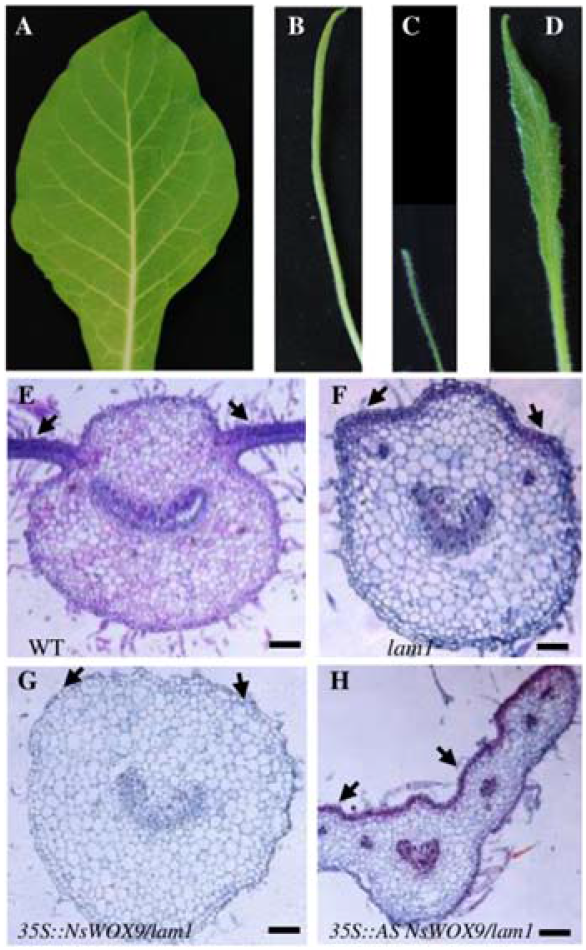
Transverse section of the leaf blade showing enhancement of the *lam1* blade by *35S::NsWOX9* and partial complementation by *35S::NsWOX9-antisense.*. **(A)** *N. sylvestris* WT leaf blade. **(B)** Leaf blade of untransformed *lam1* mutant control. **(C)** Leaf blade of *lam1* transformed with *35S::NsWOX9-antisense*. **(D)** Partially complemented leaf blade of *lam1* transformed with *35S::NsWOX9-antisense*. **(E)** Transverse section of *N. sylvestris* WT leaf blade. **(F)** Transverse section of untransformed *lam1* mutant leaf blade. **(G)** Transverse sections of *lam1* leaf blade transformed with *35S::NsWOX9* showing radialized blade. **(H)** Transverse sections of *lam1* leaf blade transformed with *35S::NsWOX9-antisense* showing blade outgrowth. Arrows indicate blade tissue in **(E)** and **(H),** vestigial blade stripes in (F) and position of blade in (G). Scale bars: 50 μm.

Transverse sections through the leaf blades showed that the *lam1* leaves had vestigial blade strips at the position of wild type blades (Figure 3F), but these strips were completely absent and blades became fully radialized in *NsWOX9* overexpressing *lam1* lines (Figure 3G). In *NsWOX9-antisense lam1* lines, on the other hand, distinct blade outgrowth was apparent but the nascent blades were not fully expanded compared to the wild type leaves (Figure 3E and 3H), confirming that blade development in the *NsWOX9-antisense* plants initiated more effectively that in the *lam1* plants but was not completed. Taken together, these results indicate that *WOX9* functions oppose those of *STF/LAM1* to negatively regulate leaf blade outgrowth in two unrelated eudicot species *M. truncatula* and *N. sylvestris*.

### *WOX9* overexpression severely affects leaf architecture

To further examine the effect of *WOX9* in leaf blade outgrowth, we introduced *35S::MtWOX9-1, 35S::MtWOX9-2* and *35S::NsWOX9* into wild type (WT) *N. sylvestris*. Analysis of over 20 independent transgenic lines for each construct revealed that all transgenic lines displayed an array of leaf phenotypes that can generally be grouped into severe and mild based on the phenotype strength. The wild type *N. sylvestris* leaf blade is a well-expanded flat lamina with smooth margin and distinctive pinnate venation pattern (Figure 4A). In plants with the severe phenotype *WOX9* overexpression completely disrupted this pattern resulting in highly distorted leaf forms. These include narrow and downward curling blades, deep margin serrations, disorganized venation patterns, uneven blade surfaces, as well as retarded plant growth with 2 to 5 tillers and additional leaves leading to a bushy appearance until stem elongation at a later developmental stage (Figures 4B, 4D, 4E, 4G; Supplemental Figure 4). While most overexpressing plants showed this strong phenotype, some exhibited a mild phenotype where the blade margin, shape and venation patterns were largely intact but with puckered and uneven blade surfaces (Figures 4C and 4F; Supplemental Figure 4). *MtWOX9-1* overexpression in *M. truncatula* also produced downward curling and narrower leaves similar to that seen in *N. sylvestris* (Supplemental Figure 5), albeit more mild, with an insignificant effect on tillering. These *WOX9* ectopic expression phenotypes in wild type and in *stf* and *lam1* mutants suggest that *WOX9* could be a negative regulator of leaf blade outgrowth antagonizing the function of *STF/LAM1*.

**Figure. 4.**
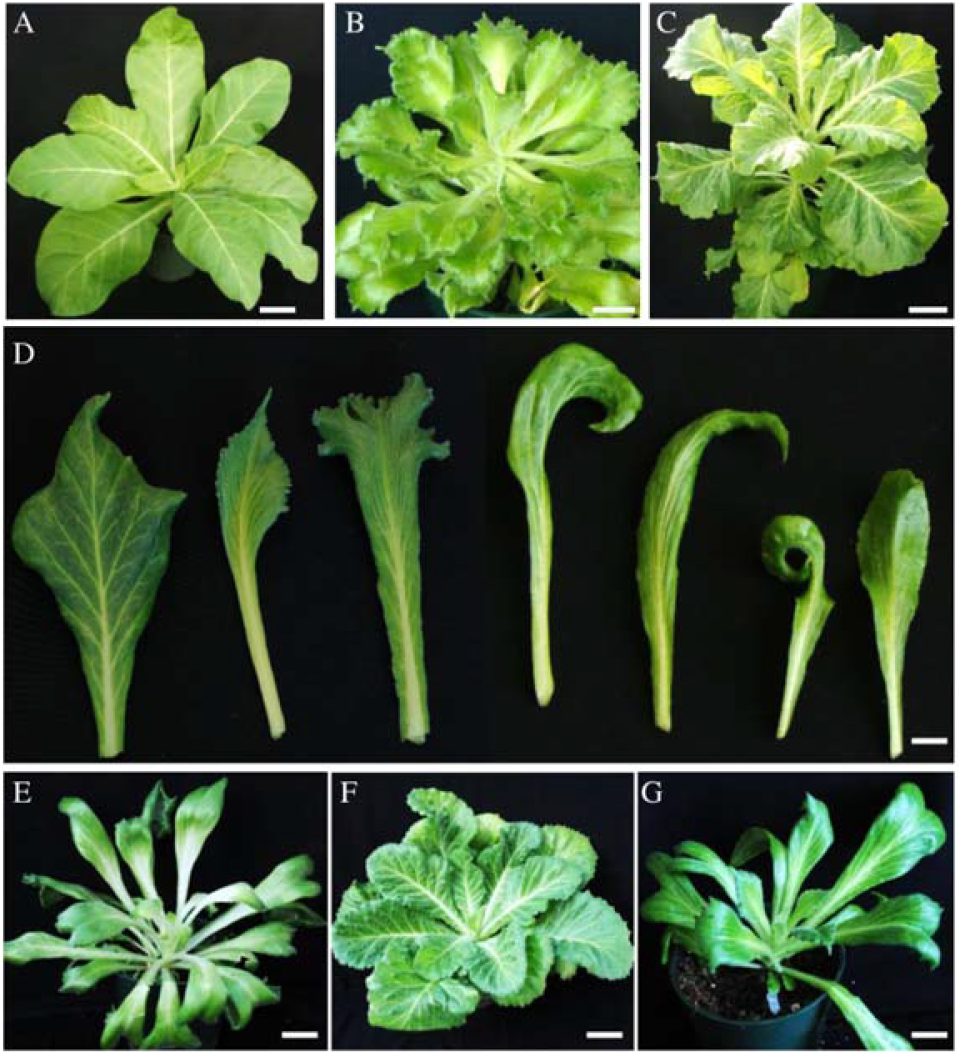
WOX9 ectopic expression in WT *N. sylvestris* alters leaf architecture. **(A)** WT *N. sylvestris* control plant at 7 weeks. **(B)** Phenotype of *35S::MtWOX9-1/WT* (severe phenotype) at 7 weeks after regeneration. **(C)** Phenotype of *35S::MtWOX9-1/WT*(mild-phenotype) at 7 weeks after regeneration. **(D)** Phenotypes of different individual representative leaves from different *35S::MtWOX9-1/WT* plants showing severe phenotypes at variable stages. **(E)** Phenotype of *35S::MtWOX9-2/WT* (severe phenotype) at early growth stage. **(F)** Phenotype of *35S::MtWOX9-2/WT* (mild) at early growth stage. **(G)** Phenotype of *35S::NsWOX9/WT* (severe) at early growth stage. Scale bars: A-C and E-G, 10 cm, D, 3 cm.

### Deleting *NsWOX9* using multiplex gRNA CRISPR/Cas9 genome editing in *N. sylvestris* alters blade symmetry and expansion

To gain insight into the function of the endogenous *WOX9* gene in wild type plants, we disrupted *NsWOX9* in *N. sylvestris* using CRISPR/Cas9 genome editing technology. We constructed an *NsWOX9*-multiplex gRNA-CRISPR/Cas9 vector containing three guide RNAs; gRNA1, gRNA2 and gRNA3 (Figure 5A), and introduced this construct into *N. sylvestris*. A total of 24 transgenic lines were generated and examined for mutations at the targeted regions. Sixteen putative mutant lines were identified by PCR amplification of the target regions using specific primers, and Sanger sequencing, indicating a 67% overall mutagenesis efficiency. Out of these sixteen putative mutants, five representative lines were selected for further characterization of their mutant phenotypes (Figure 5B). Target site sequence analysis of the five *NsWOX9*-CRISPR-mutants labeled here as *NsWOX9-1, NsWOX9-2, NsWOX9-13, NsWOX9-18 and NsWOX9-22* using SeqMan Pro 15.0.1 (DNASTAR software) revealed five different patterns of deletions ranging from 14 to 183 bp (Figure 5B). *NsWOX9-1* and *NsWOX9-13* had identical 14 bp deletions in gRNA3. *NsWOX9-2* showed two deletion events at gRNA2 and gRNA3 with 7 and 21 bp deletions, respectively. *NsWOX9-22* showed a 73 bp deletion spanning the upstream and downstream region of gRNA3, while the largest deletion was detected in line *NsWOX9-18* where a 183 bp region between gRNA1 and gRNA2 was removed, which included the PAM region of gRNA1 and extended to three nucleotides upstream of the PAM of gRNA2 (Figure 5B). All of the five CRISPR-derived mutant lines displayed malformed leaves including narrow and twisted blades, blade asymmetry, half leaf blade deletion, rough blade surface, leaf shape distortions, multiple tillers, early flowering, sterility and reduced fertility (Figures 5C to 5I). These results indicate that the negative regulation of leaf blade expansion by *NsWOX9* is required for proper leaf blade development in *N. sylvestris*.

**Figure. 5.**
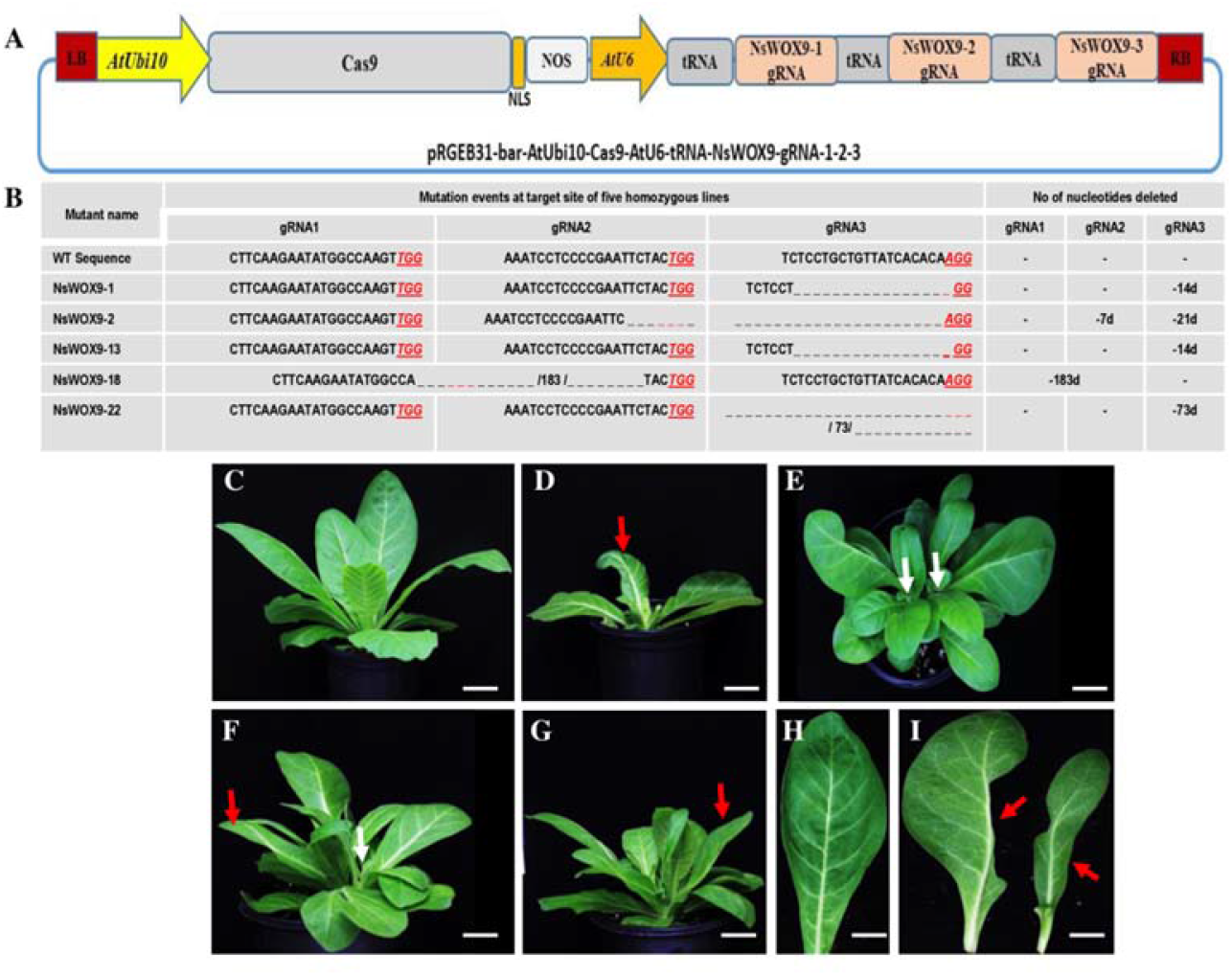
Knock out of *NsWOX9* with multiplex *gRNA-CRISPR/Cas9* in *N. sylvestris* alters leaf architecture. **(A)** Schematic representation of the three guide RNAs *NsWOX9*-gRNA1, 2 and 3 inserted in pRGEB31-bar-AtUbi10-AtU6-tRNA-gRNA vector. **(B)** Mutation events detected at the corresponding target sites of gRNA1, gRNA2 and gRNA3) in five independent CRISPR/Cas lines (*NsWOX9-1, 2, 13, 18 & 22)*. **(C-I)** Phenotype of CRISPR/Cas9 edited plants and leaves; WT control **(C)**, edited *NsWOX9-13* mutant **(D)**, edited *NsWOX9-22* mutant **(E)**, edited *NsWOX9-18* mutant **(F)**, edited *NsWOX9-2* mutant **(G)**, control WT leaf blade **(H)**, representative individual leaf blades from edited plants; left, half blade deleted (*NsWOX9-22*), and right, narrow and asymmetric blade (*NsWOX9-2*). Red arrows point to blade defects, white arrows show multiple shoots. Scale bars: A-G, 5 cm, H and I, 2.5 cm.

### *MtWOX9-1* transcript is weakly expressed in leaves and directly repressed by STF in *M. truncatula*

Reverse transcriptase quantitative PCR (RT-qPCR) analysis showed that expression of MtWOX9-1 was relatively weak in most plant tissues, including leaves, while higher levels of expression were detected in the shoot apex, flowers, pods and immature seeds, with highest levels detected in developing seeds 10 days after anthesis, followed by expression in flowers (Figure 6A). Highest expression of *MtWOX9-2*, on the other hand, was detected in the leaves followed by expression in shoot apices (Supplemental Figure 6). To examine the spatial distribution of expression, we fused a 3kb promoter region of *MtWOX9-1* upstream of the translational start codon to the β-glucuronidase (*GUS*) coding region, and transformed it into *Medicago* R108 leaf explants. GUS staining analysis revealed that expression in the mature leaf was relatively weak with particularly strong expression in the pulvinus at the base of the leaflets (Figure 6B). Strong expression was detected in the shoot apex and immature leaves, followed by flowers and stems (Figures 6C to 6E) but, no staining was detected in the anthers (Figure 6E). Very strong expression of *MtWOX9-1* was detected in immature pods and seeds at early stages of development (Figures 6F to 6I)). Interestingly, expression in the seed became progressively restricted as the seed develops, and confined only to the hilum in the mature seed (Figure 6I). These expression patterns suggest that the MtWOX9-1 function may be more important at the early stages of development, particularly during embryogenesis and leaf morphogenesis.

**Figure. 6.**
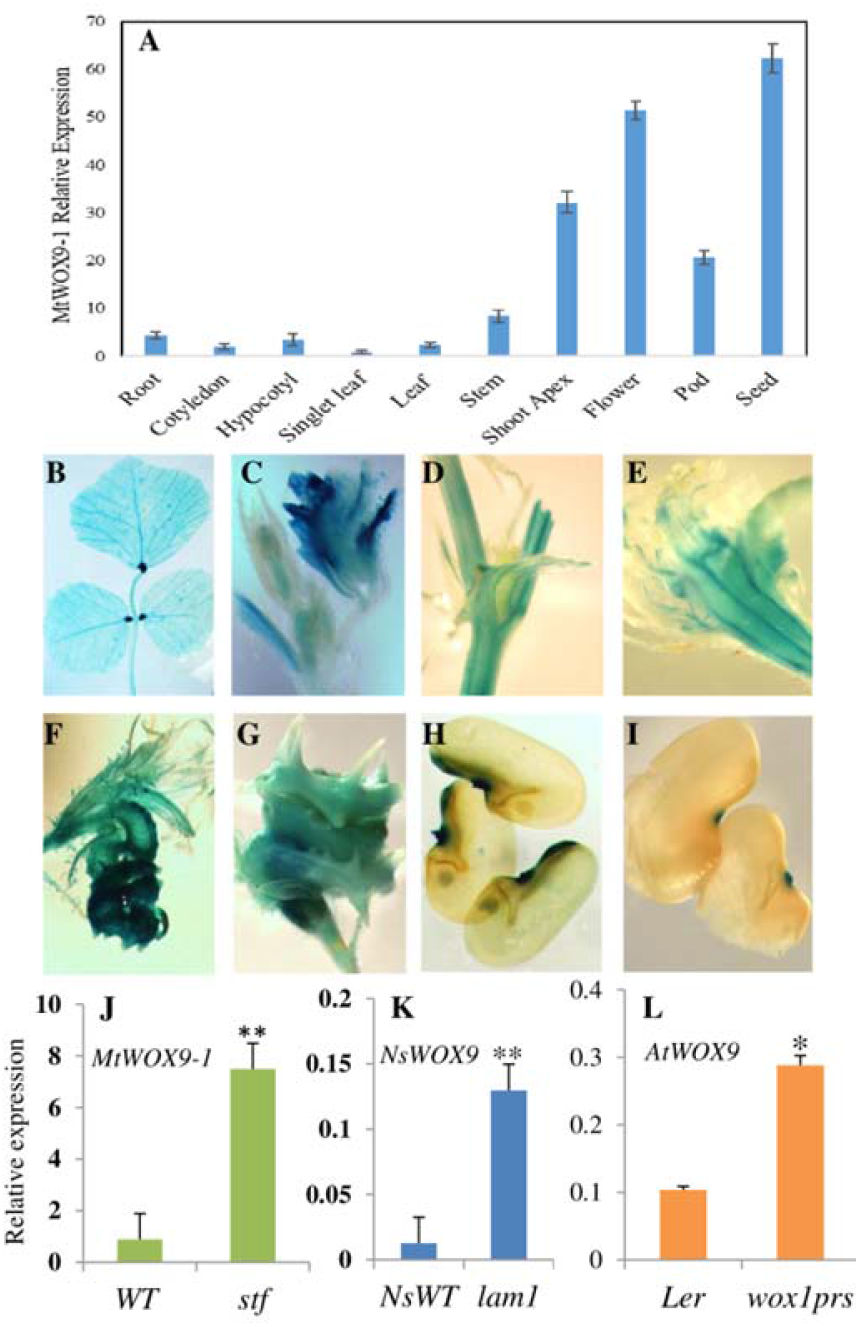
*MtWOX9-1* is weakly expressed in leaf but strongly upregulated in the *stf* mutant leaf of *M. truncatula* and corresponding mutants in *N. sylvestris* and Arabidopsis. **(A)** RT-qPCR analysis showing relative expression of *MtWOX9-1* in different tissues *M. truncatula*. leaf, stem, and shoot apex were from 4-week old plants, pods and seeds were 10 days after pollination, flower at anthesis, all the rest were at seedling stage. **(B)-(I)** GUS staining in *MtWOX9-1::GUS* transformed lines of *M. truncatula* showing fully expanded leaf **(B)**, shoot apex with folded leaves **(C)**, stem **(D)**, flower with unstained anthers **(E)**, very young pod with seeds **(F)**, older pod with seeds **(G)**, immature seeds **(H)**, and matured seeds **(I)**. **(J)** Relative expression of *MtWOX9-1* in 4-week old *stf* mutant leaf in *M. truncatula*. **(K)** Relative expression of *NsWOX9* in 4-week old *lam1* mutant leaf of *N. sylvestris*. **(L)** Relative expression of *AtWOX9* in the leaf of 4-week old Arabidopsis *wox1 prs* double mutant. Error bars indicate ± SE (n=3). Asterisks indicate significant difference from the control (*p<0.05, **p<0.01, student t-test).

To investigate the mechanistic relationship between WOX9 and positive regulators of blade outgrowth, we examined the leaf blade expression levels of *WOX9* in *stf, lam1*, and *wox1 prs*, mutants of *M. truncatula, N. sylvestris*, and *A. thaliana* respectively. RT-qPCR analyses showed that expression of *MtWOX9-1, NsWOX9*, and *AtWOX9* was upregulated by 2-4 fold in the leaves of *stf, lam1* and *wox1 prs* mutants compared to their respective wild type levels (Figures 6J to 6L), indicating that *WOX9* may be directly or indirectly repressed by the action of STF/LAM1/WOX1 in wild type leaves.

To examine whether *MtWOX9-1* is a direct target of STF, we performed a dexamethazone (DEX) induction experiment using the glucocorticoid (GR) system in the presence of the protein synthesis inhibitor, cycloheximide (CHX). Analysis was performed in 4-week old *stf* mutant plants transformed with the *35S::YFP-GR-STF* construct. The shoot apex and young leaves of the transgenic plants were treated with both DEX and CHX for 3 hours, and *MtWOX9-1* transcript accumulation was monitored by RT-qPCR in the leaves with and without the induction treatment. Our results showed that the expression of *MtWOX9-1* was reduced by approximately 60% in the DEX and CHX treated lines compared to the control CHX alone (Figure 7A). Since new protein synthesis is inhibited by CHX in the treated lines, this result suggests that *MtWOX9-1* may be directly repressed by STF. We repeated this experiment in *N. sylvestris* using *35S::YFP-GR-LAM1* fusion and found approximately 60% repression of *NsWOX9* upon *LAM1* induction by DEX plus CHX treatment (Figure 7B). However, in the reciprocal experiment, induction of *NsWOX9* expression had no significant effect on *LAM1* expression (Figure 7C), indicating that *NsWOX9* does not regulate *LAM1* transcription.

**Figure. 7.**
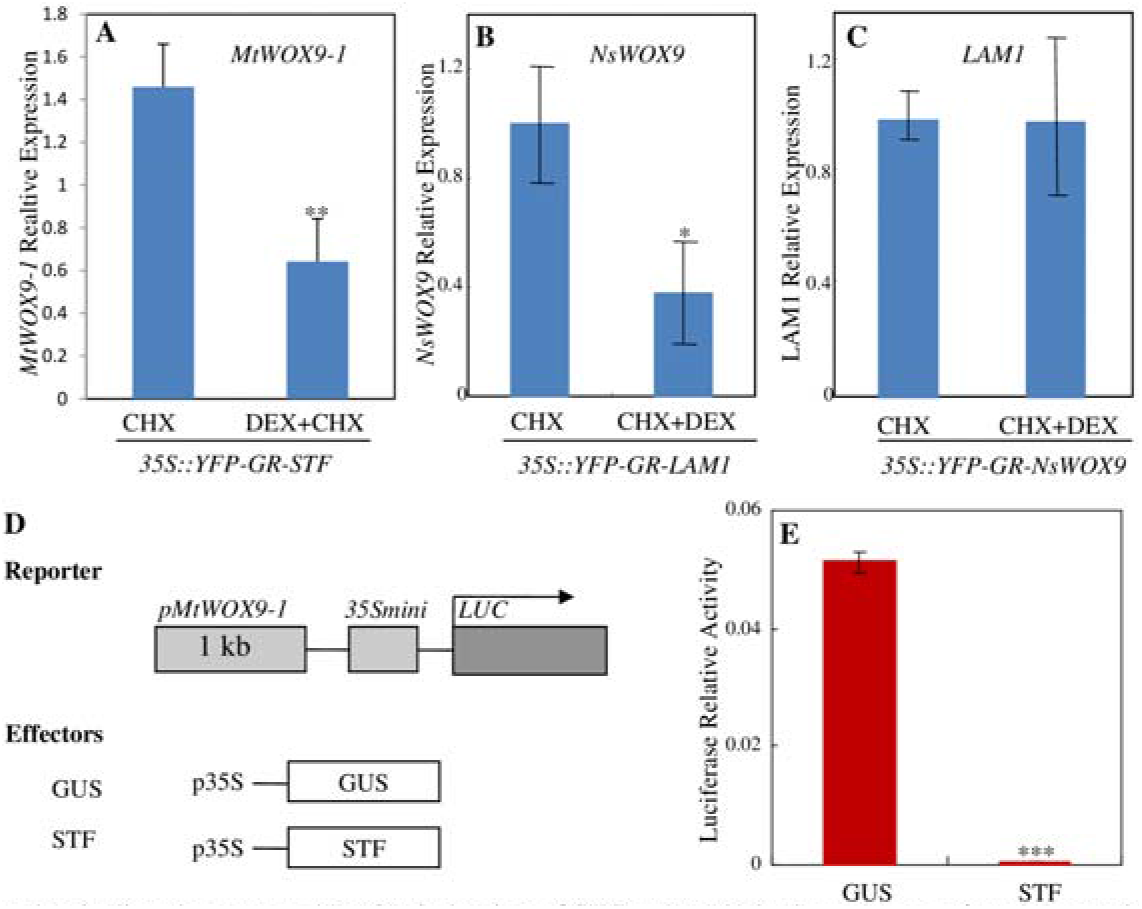
*WOX9* expression is directly repressed by GR induction of STF or LAM1 in the presence of cyclohexamide, and by STF in dual luciferase assay. **(A)** Relative expression of *MtWOX9-1* in *35S::YFP-GR-STF* transformed *M. truncatula* lines with CHX or DEX+CHX treatment for 3 hours in 4-week old leaves. **(B)** Relative expression of *NsWOX9* in 2 independently transformed *35S::YFP-GR-LAM1 N. sylvestris* lines with CHX alone, CHX+DEX or DEX alone treatments for 6 hours in 4-weeks old leaves. **(C)** Relative expression of *LAM1* in *35S-YFP-GR-NsWOX9* transformed 2 independent *N. sylvestris* lines with CHX alone, CHX+DEX or DEX alone treatments for 6 hours in 4-weeks old young leaves. Relative gene expression was determined by RT-qPCR analyses. **(D)** Schematic representation of reporter and effector constructs used in the dual luciferase assay. **(E)** Relative expression of luciferase activity (luminescence) in the presence of 35S::STF effector compared with the 35S::GUS control in Arabidopsis protoplasts. Error bars indicate ±SE (n=3). Asterisks indicate significant difference from the control (*p<0.05, **p<0.01, ***p<0.001, student t-test).

To determine if STF reduces *MtWOX9-1* transcript accumulation by directly targeting its promoter, we performed dual luciferase assay in Arabidopsis protoplasts using the Firefly-Renilla Dual-luciferase assay system (Promega). In the reporter construct, a 1kb promoter region of *MtWOX9-1* upstream of the translation start codon was fused to a mini 35S promoter driving the luciferase reporter gene (Figure 7D), while the effector constructs were made using either STF, or GUS as a negative control, both driven by the 35S promoter (Figure 7D). Consistent with the above results, co-expression of the STF effector in the protoplast almost fully abolished luciferase luminescence compared to the GUS effector control (Figure 7E), indicating that this 1 kb region of the *MtWOX9-1* promoter is sufficient for STF-dependent repression of transcription.

To determine if STF indeed binds to the *MtWOX9-1* promoter *in vitro* and *in vivo*, we performed electrophoretic mobility shift assay (EMSA) using biotin labeled probes, and chromatin immunoprecipitation (ChIP) assay, using anti GFP antibody. We previously reported that STF binds preferentially to “AT-rich” DNA elements without a strong consensus sequence (Zhang et al., 2014). We screened six such selected regions in the 3 kb upstream region of the *MtWOX9-1* promoter, and found that the MBP-STF fusion protein was able to bind to three of them (Figure 8A). These STF-binding elements are located at -22, -226 and -491 bp upstream of the *MtWOX9-1* CDS while binding was not detected with the control maltose binding protein (MBP) alone (Figure 8B). Each of these sites were significantly competed by addition of 50-fold excess of the respective unlabeled probes, indicating binding specificity. This shows that, at least *in vitro*, STF can directly bind to multiple sites within the 1 kb fragment of the *MtWOX9-1* promoter, consistent with the dual luciferase assay and GR induction experiments.

**Figure. 8.**
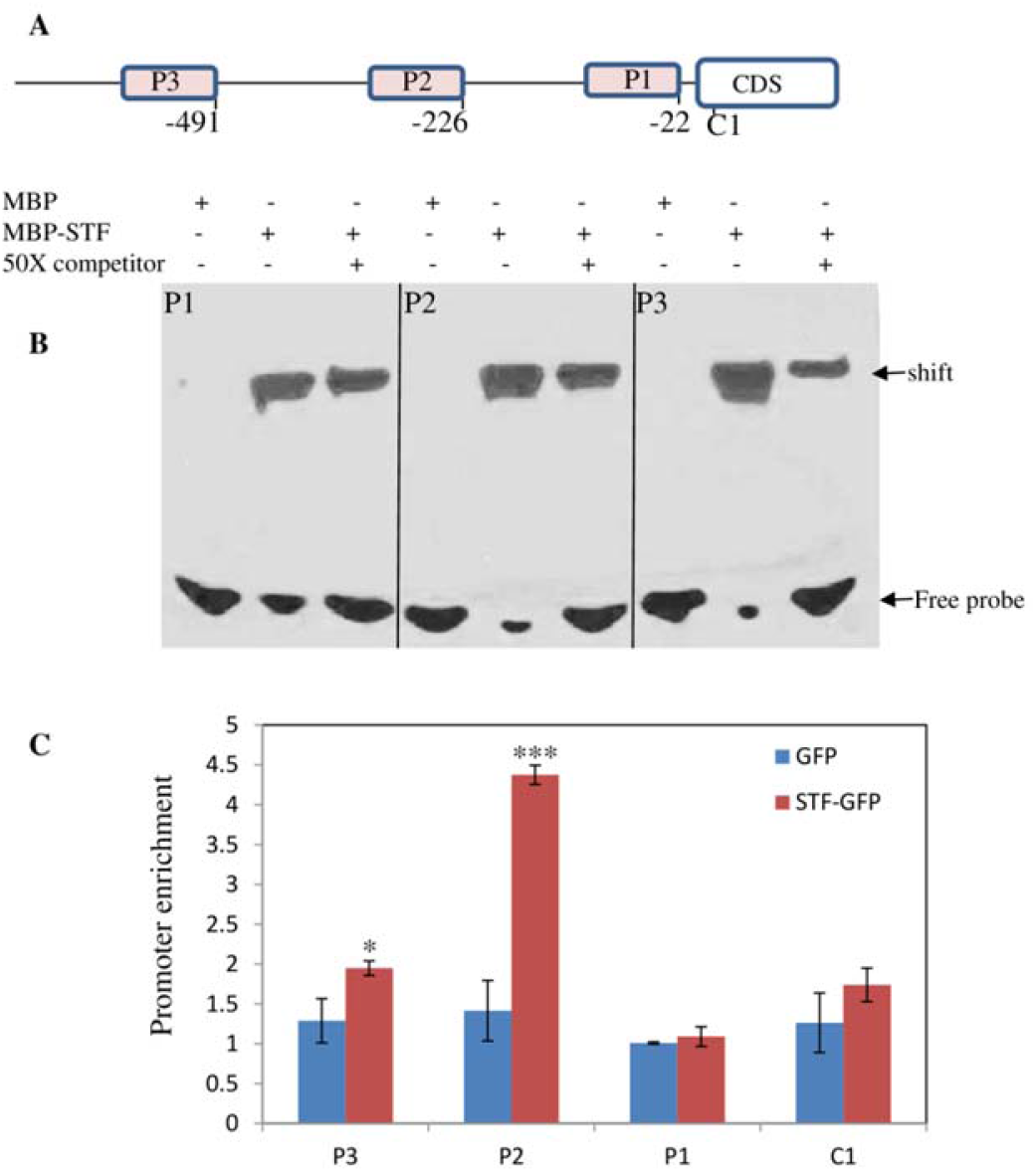
STF directly binds to the *MtWOX9-1* promoter in EMSA and ChIP assays. **(A)** Schematic representation of the *MtWOX9-1* promoter and CDS regions tested for EMSA and ChIP assays. The 3 promoter regions tested are indicated as P1, P2 and P3 and the *MtWOX9-1* coding region as C1. **(B)** EMSA showing MBP-STF bound to the biotin-labeled probe at P1, P2 and P3 promoter fragments but not the MBP control alone. Fifty-fold excess of unlabeled P1, P2 or P3 DNA was used to compete with the respective labeled probe (right lanes). **(C)** *MtWOX9-1* promoter enrichment at P1, P2, P3 regions and CDS C1. Chromatin precipitated with anti-GFP antibody from *35S::STF-GFP* and *35S::GFP* control lines were compared. Purified DNA from the chromatin were used as templates for qPCR. Note that P2 is highly enriched in *35S::STF-GFP* samples. Error bars indicate ±SE (n=3). Asterisks indicate statistical significance (*p<0.05, ***p<0.001, student t-test).

To confirm that STF binds to the *MtWOX9-1* promoter in vivo, we performed ChIP assays in leaves of Arabidopsis *wox1 prs* double mutant plants transformed with *pMtWOX9-1::MtWOX9-1* construct. Protoplasts were isolated from these transgenic plants and transformed with *35S::STF-GFP* fusion from which chromatin was isolated for analysis. ChIP-qPCR analysis revealed that among the three tested promoter regions (P1, P2 and P3), STF was highly enriched at P2 with significant enrichment also at P3 (Figure 8C). In contrast, no significant enrichment was detected at P1 or within the *MtWOX9-1* coding region (C1), indicating that STF binds strongly at the P2 position and to a limited extent at the P3 region *in planta*. Taken together, these results indicate that STF directly binds to the proximal region of the *MtWOX9-1* promoter in *M. truncatula*, and represses its activity, and that STF and WOX9 function antagonistically to regulate leaf blade outgrowth.

## Discussion

Plant specific WOX transcription factors regulate a variety of plant developmental programs from embryogenesis to shoot apical meristem maintenance and lateral organ development (Mayer et al., 1998; Schoof et al., 2000; Lohmann et al., 2001; Matsumoto and Okada, 2001; Nardmann et al., 2004; Sarkar, 2007; Breuninger et al., 2008; Shimizu et al., 2009; Vandenbussche et al., 2009; Ji et al., 2010; Tadege et al., 2011b; Nakata et al., 2012). WOX family members are also known for their promiscuous ability to substitute for each other’s functions. For example, in Arabidopsis, WUS complements the *prs/wox3* and *wox5* mutants, which are defective in floral organ development and root apical meristem maintenance, respectively (Sarkar, 2007; Shimizu et al., 2009). Conversely, members of the WUS clade WOX genes (WOX1-WOX7), with the exception of WOX4, can substitute for WUS function in stem cell maintenance (Dolzblasz et al., 2016). Arabidopsis WUS and WOX1-WOX7 can also complement the *lam1* leaf blade mutant of *N. sylvestris* (Tadege et al., 2011b; Lin et al., 2013). Here we show that the *M. truncatula* and *N. sylvestris WOX9* homologues, *MtWOX9-1, MtWOX9-2* and *NsWOX9* function antagonistically to *STF/LAM1* by negatively regulating blade outgrowth. Ectopic expression of these genes enhanced the *stf* and *lam1* leaf mutant phenotypes, and severely affected blade expansion and morphology in wild type *N. sylvestris* with a range of phenotypes (Figures 1 and 4). Conversely, reducing *NsWOX9* transcript levels in the *lam1* mutant with antisense technology partially complemented the mutant phenotype (Figure 2), indicating that *WOX9* antagonizes leaf blade outgrowth in the *STF/LAM1* pathway. However, complete knockout of *NsWOX9* by CRISPR/Cas9 genome editing technology in the wild type background resulted in a range of leaf blade deformations including lack of bilateral symmetry, altered venation patterns, narrow blades and bushy shoots (Figure 5), indicating that WOX9 function is required for proper leaf blade development.

In petunia and tomato, *WOX9* homologues are involved in inflorescence development and architecture (Lippman et al., 2008; Rebocho et al., 2008; Costanzo E, 2014). Both the *evergreen (evg*) mutant in petunia (Rebocho et al., 2008) and *compound inflorescence* (*s*) in tomato (Lippman et al., 2008), which dramatically alter the wild type inflorescence architecture are caused by lesions in *WOX9* homologues. The *s* allele in tomato results in a highly branched structure with hundreds of flowers, which increases fruit production and may have been selected by breeders ((Lippman et al., 2008). In the *evg* mutant of petunia, on the other hand, the inflorescence stem often fails to bifurcate after the formation of bracts and continues to growth as a single thickened stem without physical separation of the floral meristem (FM) and inflorescence meristem (IM), leading to a fasciated appearance (Rebocho et al., 2008). Unlike *s*, the *evg* FM also fails to produce floral organs suggesting that EVG is required for inflorescence bifurcation and floral organ identity, though *evg* mutants are indistinguishable from wild type during early vegetative growth (Rebocho et al., 2008). Thus, in tomato and petunia, *WOX9* homologues appear to have opposite effects specific to inflorescence development and architecture. However, both tomato and petunia have a cymose inflorescence pattern (determinate growth) and it is unclear whether these inflorescence-associated defects are specific to cymose or are also exhibited by racemose (indeterminate growth) and panicle (mixed inflorescence) inflorescences. At least in Arabidopsis (racemose inflorescence), the role of WOX9 appears not to be restricted to inflorescence development, and in rice (mixed inflorescence), WOX9 is involved in uniform tiller growth and development (Wang et al., 2014; Fang et al., 2020).

In Arabidopsis, *WOX9*, also called *STIMPY (STIP*), is required for meristem growth and maintenance and positively regulates *WUS* (Wu et al., 2005). *stip* mutants display arrested growth at an early stage of development but can be fully rescued by sucrose (Wu et al., 2005). STIP/WOX9 is shown to mediate cytokinin signaling during shoot meristem establishment and, together with WOX2 and WOX8, regulates zygote and embryo polarity patterning (Wu et al., 2007; Breuninger et al., 2008; Skylar et al., 2010; Ueda et al., 2011). WOX9 homologues in other species are also reported to be involved in promoting somatic embryogenesis (Gambino et al., 2011; Tvorogova, 2019). In all of these examples, the function of WOX9 appears to center on cell proliferation and/or meristematic competence for proper plant growth and development. Our observation of the effect of WOX9 overexpression and knockout in Medicago and woodland tobacco leaf development is consistent with these findings, and may reflect a conserved molecular function in cell proliferation and differentiation during growth and development. For example, the Arabidopsis gain-of-function mutant *stip*-D displays wavy leaf margins and increased number of axillary shoots leading to a bushy phenotype (Wu et al., 2005). A phenotype similar the wavy margins and bushy shoots seen in all *MtWOX9-1, MtWOX9-2* and *NsWOX9* overexpressing *N. sylvestris* transgenic lines (Figure 4), which suggests misregulation of cell proliferation in leaf primordia. The observation that both overexpression and knockout of *WOX9* in *N. sylvestris* led to narrow leaf and bushy shoot phenotypes suggests that WOX9 may be involved in maintaining a balance between cell proliferation and differentiation, necessitating that *WOX9* transcripts be maintained at a required optimum. However, the mechanism of WOX9 function and control of its steady state transcript levels during leaf morphogenesis and maturation is not well understood. We previously reported AtWOX9 to be a transcriptional activator, based on its unique effects on the *lam1* mutant of *N. sylvestris* (Lin et al., 2013), which could provide insight into its molecular function.

The Arabidopsis WOX family has been divided into three clades based on phylogenetic analysis: the modern/WUS clade (WUS, WOX1-7), intermediate clade (WOX8, 9, 11, 12), and ancient clade (WOX10, 13, 14) (van der Graaff et al., 2009). This classification is largely consistent in other species as well (Zhang et al., 2010; Hao et al., 2019; Wu, 2020). The WUS clade members are characterized by an intact WUS box motif (Haecker et al., 2004; Lin et al., 2013), which is a transcriptional repression motif (Ikeda et al., 2009; Lin et al., 2013). Members of this group function primarily as transcriptional repressors, able to complement the *lam1* mutant phenotype (Lin et al., 2013), and are capable of substituting for WUS function in maintaining vegetative and floral meristems (Dolzblasz et al., 2016). The WOX1 homologues *M. truncatula* STF and *N. sylvestris* LAM1 belong to this clade and function as master regulators of leaf blade outgrowth through a transcriptional repression mechanism in association with the co-repressor TOPLESS (Tadege et al., 2011b; Lin, 2013; Lin et al., 2013; Zhang et al., 2014; Zhang et al., 2019). The intermediate and ancient clade members have partial or no WUS box, and do not have transcriptional repression activity in dual luciferase assays. As a result, they are unable to rescue the *lam1* mutant (Lin et al., 2013) nor substitute for WUS function (Dolzblasz et al., 2016). Among the intermediate and ancient clades, AtWOX9 is unique in that it displays the strongest activation activity in dual luciferase assays, and strongly enhances the *lam1* mutant phenotype, affecting blade outgrowth in both medial-lateral and proximal-distal axes (Lin et al., 2013), indicating that transcriptional activation activity modulated by AtWOX9 is antagonistic to LAM1 function. This is consistent with the observation that activation activity at the *STF* expression domain antagonizes *STF* function in blade outgrowth (Zhang et al., 2014; Zhang et al., 2019).

The results presented here demonstrated that *WOX9* transcript is upregulated in three leaf blade mutants; *stf* in *M. truncatula, lam1* in *N. sylvestris* and *wox1 prs* in Arabidopsis (Figures 6J to 6L), indicating that *WOX9* transcription may be suppressed by STF/LAM1/WOX1 in these species to allow blade outgrowth. Several lines of evidence including a GR inducible system in the presence of DEX and CHX, dual luciferase assay, EMSA, and ChIP confirmed that STF/LAM1 directly binds to the *MtWOX9-1* promoter to repress *WOX9* transcription (Figures 7 and 8), demonstrating that modern clade WOX1/STF/LAM1-mediated repression of intermediate clade *WOX9* is required for proper leaf blade outgrowth in eudicots. In Arabidopsis, *WOX9* functions upstream of *WUS* and is supposed to activate *WUS* to promote vegetative meristem growth (Wu et al., 2005), although it is unclear whether this activation is direct or indirect. *WOX9* is activated by cytokinin signaling (Skylar et al., 2010), and type-B ARRs directly activate *WUS* (Meng et al., 2017; Wang et al., 2017; Zhang et al., 2017; Zubo et al., 2017; Xie, 2018), while WUS promotes cytokinin activity by repressing type-A ARRs (Leibfried et al., 2005). Thus, cytokinin signaling provides a potential connection between WOX9 and WUS in Arabidopsis, but whether WUS can directly affect WOX9 activity via negative or positive feedback loop is yet to be determined. Our work clearly demonstrates that in Medicago and woodland tobacco, the WUS clade member STF/LAM1 directly represses *MtWOX9-1* or *NsWOX9*, but significant *STF/LAM1* activation by WOX9 was not detected (Figure 7C). Although MtWOX9-1 may not activate *STF*, it is likely to activate other targets in leaf development. WOX9 amino acid sequences from different species show a highly conserved acidic domain at the C-terminus (Supplemental Figure 1), which could mediate transcriptional activation. Thus, our current model is that STF represses the transcription of key targets at the adaxial-abaxial junction to promote cell proliferation, and these targets include *MtWOX9-1* and cell differentiation factors such as *AS2* (Zhang et al., 2014). WOX9, on the other hand, may negatively regulate blade outgrowth by directly activating targets independent of STF and/or by activating targets repressed by STF (Figure 9). The hypothesis that STF and WOX9 may oppositely regulate common targets providing a critical balance between cell proliferation and differentiation during leaf morphogenesis, explains why WOX9 ectopic expression enhances the *stf/lam1* mutant phenotype. Since both STF and WUS promote cell proliferation via a transcriptional repression mechanism in the leaf primordium and SAM, respectively, our findings suggest that direct control of *WOX9* activity by WUS may be required for SAM maintenance as well, uncovering a mechanistic framework for WOX modulated control of robust plant growth and developmental programs. It would be interesting to investigate if repression of the intermediate and ancient clade WOX transcriptional activators by modern clade WOX transcriptional repressors is a universal strategy exploited during the evolution of land plants and resulting in the complex morphological architecture of higher plants.

**Figure. 9.**
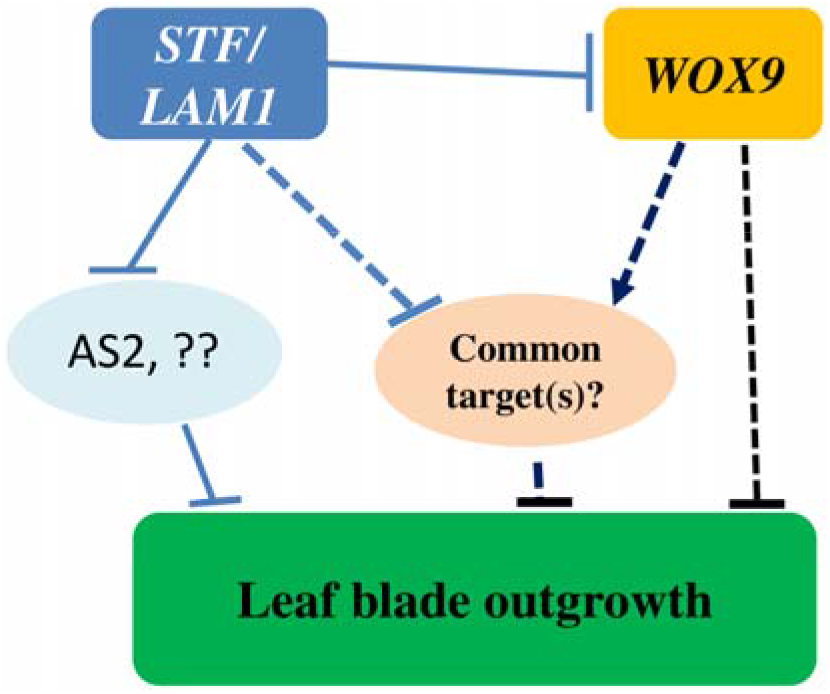
Schematic representation of hypothetical model for the regulation of leaf blade outgrowth by the interaction of STF/LAM1 and WOX9 STF/LAM1 directly represses *WOX9, AS2*, and other unidentified factors to promote leaf blade outgrowth. WOX9, on the other hand, negatively regulates leaf blade outgrowth by activating negative regulators of leaf growth and/or directly repressing blade outgrowth processes. The model proposes that STF/LAM1 and WOX9 may have a common target(s) repressed by STF/LAM1 and activated by WOX9 to balance cell proliferation with differentiation during leaf morphogenesis.

## Methods

### Plant materials and growth conditions

Plant materials used for this study including *Medicago truncatula* R108, *stf* mutant, and *Nicotiana sylvestris* (woodland tobacco) wild type and *lam1* mutant were grown in one gallon pots in a greenhouse under long-day (LD) conditions with 16/8 hours light/dark cycle at 23-27°C, and in growth room under long-day conditions of 16/8 hours light /dark cycle at 23-25°C, 70-80% relative humidity, and a light intensity of 150 μmol.m^2^.

### Samples collection, RNA extraction

*M. truncatula* tissue samples (young leaf and shoot) were collected from 4 week old wild type and *stf* mutant plants. For GR induction, four weeks old plants of *p35S::YFP-GR-STF, p35S::YFP-GR-LAM1* and *p35S::YFP-GR-NsWOX9* transgenic lines were treated with mock, DEX (10 mM), and CHX (10 mM) for 3hrs in *M. truncatula* and 6 hrs in *N. sylvestris*, before leaf samples were collected. All leaf samples from wild type, mutants and transgenic lines were collected at appropriate times for analyses of gene expression patterns as indicated in figure legends. Collected samples were immediately snap-frozen in liquid nitrogen and stored at −80°C until processing.

Total RNA from shoot apex and young leaf of *M. truncatula* R108 (WT) and *stf* mutant, *N. sylvestris* (WT) and *lam1* mutant, Arabidopsis L*er* (WT), *wox1/prs* mutant and *p35S::YFP-GR-STF, p35S::YFP-GR-LAM1* and *p35S::YFP-GR-NsWOX9* transgenic lines were isolated using TRIzol Reagent (Invitrogen) for cDNA synthesis.

### Real time PCR

Expression patterns analyses were performed using quantitative real time PCR (RT-qPCR) and semi-quantitative PCR with specific forward and reverse primers (Supplemental table 1). Reverse transcription (RT) was performed using RNA treated with DNase I (Invitrogen), an oligo (dT) primer, and SuperScript III reverse transcriptase (Invitrogen) according to the manufacturer’s instruction. Quantitative RT-PCR assays were performed with three biological repeats and three technical replications of each experiment using SYBR Green real-time PCR Master Mix (Invitrogen). *M. truncatula*, Arabidopsis and *N. sylvestris* actin primers were used as expression standards. All specific forward and reverse primers used for gene cloning and expression and related molecular analyses in this study are listed in Supplemental Table 1.

### Gene isolation and transgene construction

*MtWOX9-1* and *MtWOX9-2* genes were isolated from the *M. truncatula* genome by BLAST search using the AtWOX9 sequence. Full-length *MtWOX9-1&2* coding sequences were amplified by RT–PCR using total RNA extracted from leaf samples. To generate transgenic plants of *M. truncatula*, the cDNA was sub-cloned into the pDONR207 entry vector (Invitrogen) by BP clonase reaction. The final constructs were produced by an LR clonase reaction between each of the entry vectors and pMDC32 destination vector. The resulting plasmids were transferred into *Agrobacterium tumefaciens* strain *AGL1* and used to transform into *M. truncatula* R108 and *stf* mutant via agro-mediated-transformation using leaf explants as described (Tadege et al., 2011b). *N. sylvestris* WT and *lam1* mutant plants were transformed using *Agarobacterium tumefaciens, GV2260* strain by leaf disc inoculation method as previously described (Tadege et al., 2011b). Antisense of *N. sylvestris (NsWOX9-anti*) construct was made using the Gateway system (Invitrogen). Briefly, *NsWOX9-anti-F* and *NsWOX9-anti-R* primers were designed by reverse attachment attb2 and attb1 to the primers for amplification of the full-length CDS sequence from start and stop codon of the gene. Using such primers, *NsWOX9* was amplified from *NsWOX9-cDNA* and cloned into *pDONR207* entry vector (Invitrogen) by BP clonase reaction. Then, the construct was transferred into destination vector pMDC32 by an LR clonase reaction. All generated transgenic lines were confirmed by PCR using specific primers, and reduced expression lines were identified by RT-PCR.

### Multiplex gRNA-CRISPR/Cas9 construction

To generate the multiplex gRNA-CRISPR/Cas9 *NsWOX9* construct, we designed three multiplex gRNAs targeting multiple sites of exon 2 (gRNA1-CTTCAAGAATATGGCCA AGT; gRNA2-CTTCAAGAATATGGCCAAGT and gRNA3-TCTCCTGCTGTTATCA CACA) at the upstream of PAM (TGG, TGG, and AGG) sites, respectively, using the web-based tool CRISPR-P (http://cbi.hzau.edu.cn/cgi-bin/CRISPR) (Lei, 2014). All designed gRNAs were inserted between tRNA and gRNA scaffolds and clustered in tandem using the Golden Gate assembly method (Engler, 2008). The pGTR plasmid, which contains a tRNA-gRNA fragment, was used as a template to synthesize polycistronic tRNA-gRNA (PTG) (Xie, 2015). The overlapping PCR products were separated and purified by the Spin Column PCR Product Purification Kit (Wizard SV Gel and PCR Clean-Up System) following manufacturer’s instruction (Promega, WI). Then, the chain of multiplex tRNA-gRNA with three *NsWOX9*-spacers was inserted into an optimized vector with AtU6-tRNA-gRNAs-AtUbi10-Cas9-pRGEB31-bar backbone by digestion and ligation using *Fok I* (NEB) and *BsaI* enzymes (Xie, 2015).

### CRISPR/Cas plant transformation and analysis

The subsequent multiplex *NsWOX9spacer*-gRNA-CRISPR/Cas9 binary vector construct was transformed into *Agrobacterium* strain *GV2260*. First, transgenic lines were screened by PCR using genomic DNA and specific primers (PPT-F + PPT-R) of Barsta selection marker. Then, putative *nswox9-CRISPR* mutants were identified through amplification of the target region by PCR using extracted genomic DNA as a template with specific primers designed from the border of the target site. Amplified fragments of the mutated region were sub-cloned into pGEM-T easy plasmid by TA-Cloning and 10 colonies for each locus were subjected to Sanger sequencing. Reads were analyzed by aligning with the reference sequence using the SeqMan Pro 15.0.1 (DNASTAR software for life scientists) (https://www.dnastar.com/quote-request/).

### Electrophoretic mobility shift assay

The *MtWOX9-1* promoter with 37, 30 and 32 bp oligonucleotides, corresponding to starting sites at -22, -226 and -491, respectively, upstream of the start codon were labeled using the Biotin 3’ End DNA Labeling Kit according to the manufacturer’s instructions (Thermo Scientific/Pierce). EMSA was performed with the Light Shift Chemiluminescent EMSA Kit (Thermo Scientific/Pierce). Unlabeled probes with a 50-fold higher concentration were used as competitors in each of the competing assays. Purified maltose binding protein (MBP) and MBP-STF were used in the EMSA as described (Zhang et al., 2014). Gel electrophoresis was performed on 10% native polyacrylamide gels. After blotting on a positively charged nylon membrane (Amersham), the DNA was crosslinked using a transilluminator equipped with 312 nm bulbs with the membrane face down for 15 min. The biotin-labeled DNA was detected by Chemiluminescence and exposed to X-ray film (Kodak). The probes and primers used in EMSA assay are listed in Supplemental Table 2.

### ChIP Assays

ChIP assays were performed as described previously (Xiong et al., 2013; Chen et al., 2018). Protoplast extracted from 14-d-old *pMtWOX9-1:MtWOX9-1/wox1prs* transgenic Arabidopsis leaves were transformed with 10 μg of *35S::STF-YFP* using the polyethylene glycol–mediated transformation method. Protoplasts were cross-linked by 1% formaldehyde in W5 medium for 20 min and quenched with Glycine (0.2 M) for 5min. The protoplasts were then lysed, and the DNA was sheared on ice with sonication. The sheared chromatin was precleared by salmon sperm–sheared DNA/protein A agarose beads. Precleared chromatins were incubated with 5 μl of anti-GFP antibody (ab290) overnight at 4°C, after which Protein A agarose beads (40 μl) were added, and the samples were incubated at 4°C for 2 hrs. After reversing the crosslinks with Proteinase K at 65°C overnight, DNA was purified and analyzed by q-PCR amplification using specific primers. The input DNA and HA antibody–precipitated DNA were used as PCR templates for the positive and negative controls, respectively. Experiments were repeated three times. The primers used for the ChIP assays are listed in Supplemental Table 1

### Dual luciferase assay

For effector plasmids, the coding sequence of *STF* or GUS was cloned into pDONR207 entry vector and then transferred into p2GW7 using the Gateway system (Invitrogen). Construction of the reporter *pMtWOX9-1-mini-35S-LUC* plasmid containing 1kb of the *MtWOX9-1* promoter was performed as previously described (Zhang et al., 2014). Transient expression assays were performed in Arabidopsis protoplasts as described (Asai et al., 2002). For normalization, 0.5 μg of plasmid pRLC was used as an internal control.

### Histological analysis

Leaf samples were fixed in formaldehyde for 48 hrs and dehydrated in an ethanol series (60, 70, 85 and 95%). Then, leaves were embedded in Paraplast (Sigma-Aldrich, St. Louis, MO) and tissue sections (15 μm thick) were cut with a Reichert-Jung 2050 microtome. Specimens were mounted on slides and stained with Safranine O and Light Green as previously described (Tadege et al., 2011b). Images were captured with digital camera mounted on an Olympus BX-51 compound microscope.

### Sequence alignment and phylogenetic tree construction

Multiple protein sequence alignment was performed using BioEdit software and the ClustalW program (http://www.mbio.ncsu.edu/bioedit/bioedit.html). Species refer to At (*Arabidopsis thaliana*), Ns (*Nicotiana sylvestris*), Pc (*Phaseolus coccineus*), Gm (*Glycine max*), Vv (*Vitis vinifera*), Ph (*Petunia x hybrid*), Cs (*Cucumis sativus*), Sl (*Solanum lycopersicum*) and Ca (*Capsicaum annuum*). A neighbor-joining phylogenic tree was constructed using MEGA-X default settings with 1000 bootstrap replications (http://www.megasoftware.net/).

### Statistical Analysis

For statistical analysis, Student’s *t* test was used as specified in figure legends. Asterisks indicate statistical differences (*P < 0.05, **P < 0.01, ***p < 0.001).

### Accession Numbers

Sequence data from this article can be found in the NCBI (http://www.ncbi.nlm.nih.gov/) databases, *M. truncatula* Genome Database (http://www.medicagogenome.org/), or Phytozome (https://phytozome.jgi.doe.gov/pz/portal.html) under the following accession numbers: MtWOX9-1, Medtr2g015000; MtWOX9-2, Mt7go26130; AtWOX9, AT2G33880; AtWOX8, AT5G45980; NsWOX9, XM_009794999; Ns-LAM1, AEL30893; MtSTF, JF276252; AtWOX1, AT3G18010; AtWOX3, AT2G28610; PcWOX9, ACL11801; GmWOX9, XP_006594207; GmWOX9, XP_003541514; VvWOX9, XP_002273188; CaWOX9, XP_016562050; CsWOX9, XP_004134676; PhWOX9 (evergreen(EG)), ABO93066; PhWOX9 (SOEG); ABO93067; SlWOX9 (COI), NP_001234072; SlWOX9 (COI) isoform, XP_010315848.

## Supplemental Data

**Supplemental Figure 1.** Amino acid sequence alignment of MtWOX9-1/2 and other related eudicot WOX9 sequences.

**Supplemental Figure 2.** Phylogenetic analysis of MtWOX9-1/2 and other related eudicots WOX9 protein sequences.

**Supplemental Figure 3.** Ectopic expression of *MtWOX9-1* enhances the *lam1* mutant phenotypes.

**Supplemental Figure 4.** Leaf phenotypes of *NsWOX9* overexpression in *N. sylvestris*.

**Supplemental Figure 5.** Phenotype of *MtWOX9-1* overexpression in *M. truncatula*.

**Supplemental Figure 6.** RT-qPCR analysis of *MtWOX9-2* expression in different tissues of *M. truncatula*.

**Supplemental Table 1.** List of primers and gRNAs used in this study

## Acknowledgements

This work was supported by the National Science Foundation (NSF) grant IOS-1354422, Agriculture and Food Research Initiative Grant No. 2015-67014-22888 from the USDA National Institute of Food and Agriculture, and the Agricultural Experiment Station. Work in VT’s laboratory at St Petersburg State University, Russia, was supported by the Russian Science Foundation project no. 16-16-10011 and by grant from the Russian Foundation for Basic Research no. 20-016-00124.

## Author Contributions

T.W.W., H.W., and M.T. designed the research. T.W.W., H.W., D.T., F.Z., M.B., H.A., Y.L., and N.C. performed the experiments. T.W.W., H.W., F.Z., J.C., R.A., and M.T. analyzed the data. T.W.W., R.A., and M.T. wrote the manuscript.

